# Sir2 is required for the quiescence-specific condensed three-dimensional chromatin structure of rDNA

**DOI:** 10.1101/2024.12.12.628092

**Authors:** Christine Cucinotta, Rachel Dell, Kris Alavattam, Toshio Tsukiyama

## Abstract

Quiescence in *Saccharomyces cerevisiae* is a reversible G_0_ crucial for long-term survival under nutrient-deprived conditions. During quiescence, the genome is hypoacetylated and chromatin undergoes significant compaction. However, the 3D structure of the ribosomal DNA (rDNA) locus in this state is not well understood. Here, we report that the rDNA locus in quiescent cells forms a distinct condensed loop-like structure, different from structures observed during the mitotic cell cycle. Deletion of *SIR2* disrupts this structure, causing it to collapse into a small dot and resulting in quiescence entry and exit defects. In contrast, Sir2 affects rDNA structure only modestly in G2/M phase. In the absence of Sir2, occupancy of both RNA Polymerase II and histone H3 increase at the rDNA locus during quiescence and through quiescence exit, further indicating gross defects in chromatin structure. Together, these results uncover a previously undescribed rDNA chromatin structure specific to quiescent cells and underscore the importance of Sir2 in facilitating the transition between cellular states.

## INTRODUCTION

Quiescence, a reversible state of cell cycle withdrawal (G0), is a fundamental biological process that allows cells to survive under adverse conditions and resume proliferation upon favorable conditions^1^. This state is not only important for survival of single-celled organisms like *Saccharomyces cerevisiae*^2,3^, but also plays a crucial role in metazoans. Quiescence enables stem cells to maintain tissue homeostasis, supports T-cell activation in immune responses, and contributes to wound healing and regeneration^4,5^. Understanding the molecular mechanisms governing quiescence is therefore essential to elucidate how cells regulate their ability to exit and re-enter the cell cycle after prolonged cell cycle withdrawal.

A defining feature of quiescent cells is their highly repressive chromatin landscape, which is conserved across species^6^. This chromatin repression involves widespread histone deacetylation, narrower nucleosome depleted regions (NDRs), and an overall reduction in transcription^7–9^. Yeast cells present a dramatic difference in chromatin structure between quiescence and the cell cycle^10–13^. Such extreme chromatin reorganization makes quiescent yeast cells an excellent model to study how 3D chromatin structures are established, maintained, and their biological functions. Recent work from our lab and others has revealed that large-scale changes in chromatin structure accompany quiescence entry. Our previous work has focused on chromatin dynamics during quiescence at two levels. First, we explored changes in chromatin domains and their spatial organization^12^, followed by detailed investigations into how nucleosome arrays fold into compact structures during quiescence^13^. A larger scale 3D structural change also occurs, where telomeres in quiescent yeast cells undergo hyper-clustering, forming one or two large foci instead of multiple smaller clusters typically seen in cycling cells^14,15^. Together, these studies revealed a unique 3D chromatin landscape in quiescent cells that is distinct from actively cycling cells. However, not all the genome’s large-scale 3D structural changes have been defined in quiescence. In addition, the molecular mechanisms that control these structures and their significance in quiescence regulation remain largely unknown.

In this study, we focused on the ribosomal DNA (rDNA) locus, which in yeast is localized on the chromosome XII in ∼150 tandem repeats, occupying about 1.4 mega base pairs^16^. The rDNA locus, containing sites for all three RNA polymerases and the Fob1 replication fork block, requires complex regulation to maintain copy number and genome integrity^17^. Using super-resolution microscopy and Micro-C data analyses, we reveal that the rDNA forms a novel 3D structure specific to quiescence. We demonstrate that this structure depends on Sir2, a histone deacetylase that is well-known for its role in chromatin silencing at the rDNA and telomeres during the cell cycle^18^. We find that deletion of the *SIR2* gene causes strong defects in both quiescence entry and exit. In addition, *sir2*Δ cells have abnormally high levels of nucleosome and RNA polymerase II occupancy at the rDNA locus in quiescence as well as throughout the quiescence exit process. Together, our results provide new insights into how 3D chromatin architecture contributes to the maintenance of quiescence and the regulatory mechanisms underlying long-term cell cycle withdrawal.

## RESULTS

### rDNA in quiescent cells forms a distinct structure

To microscopically examine 3D chromatin structure in budding yeast quiescent cells, we have employed stimulated emission depletion (STED) microscopy^19^. For visualizing total chromatin, we used an antibody against H2B (Fig. 1A, top row). As expected, chromatin was much more condensed in quiescent cells compared to G_1_ and G_2_/M cells^20,21^. We unexpectedly noticed the presence of condensed loop-like structures in quiescent cell chromatin (Fig 1A right, arrows), which were smaller than those in G_2_/M cells (Fig 1A center, arrows). In contrast, loop-like structures were not found in G_1_ cells (Fig 1A left, arrows). It has been established that the rDNA locus in yeast forms loops in G_2_/M phase^22,23^. We therefore wondered whether the loop-like structures in quiescent cells are also formed by the rDNA locus. To test this possibility, we fused a Flag epitope tag to the protein Net1. Net1, a part of the RENT complex, recruits Sir2 for rDNA-specific histone deacetylation^24^, and is frequently used to visualize the rDNA locus in yeast^23^. Using an anti-Flag antibody to visualize C-terminally Flag-tagged Net1, we observed that rDNA was clearly visible across all cell states tested, including in quiescent cells (Fig. 1A, middle row). In cells in the mitotic cell cycle, we observed G_1_ rDNA “puffs” and more clearly discerned G_2_/M loops. Quiescent cells formed novel, condensed loop-like structures that have not been observed in any other cell states (see merged images in Fig. 1A, bottom row).

**Figure 1.**
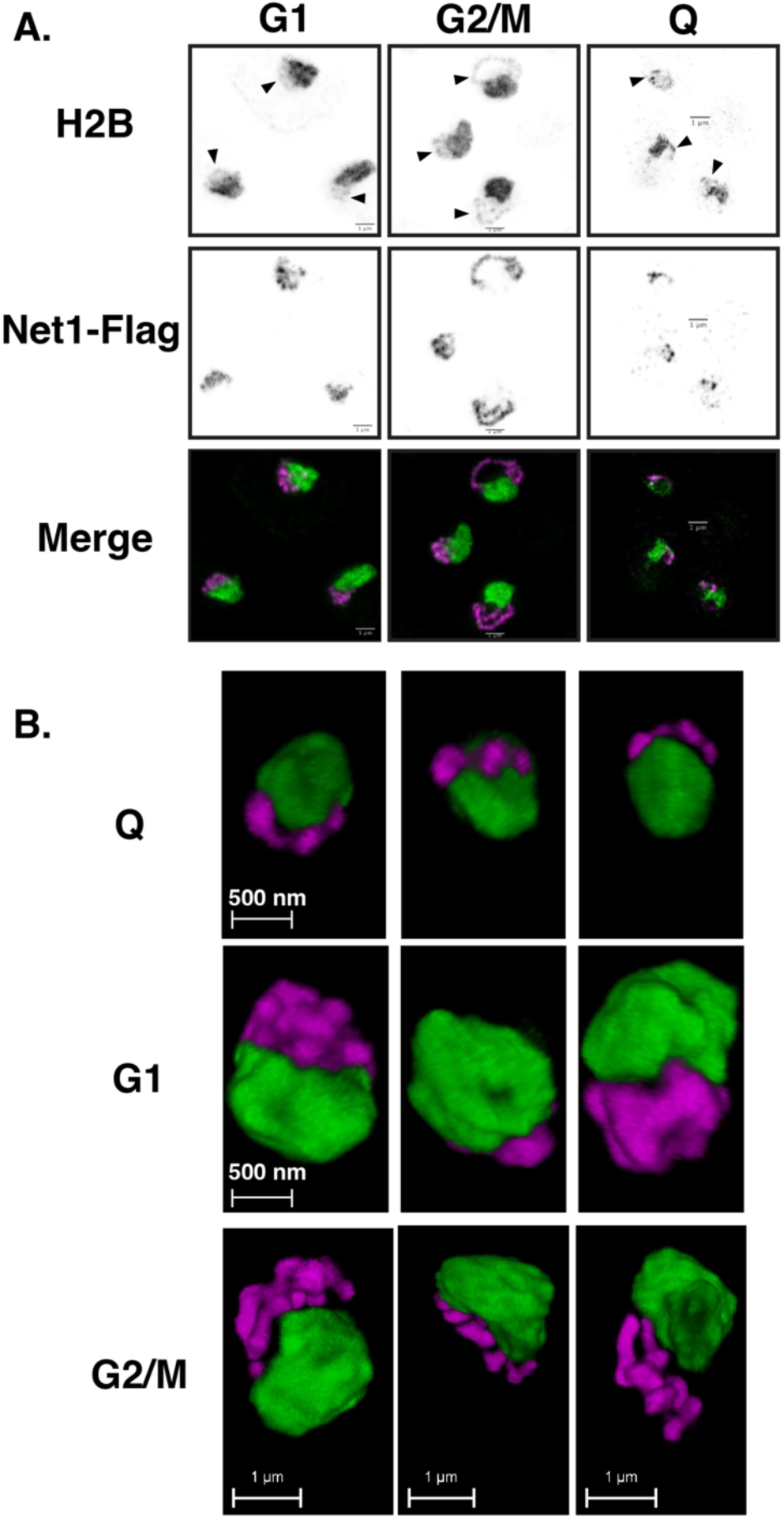
The ribosomal DNA locus (rDNA) forms a distinct structure in quiescent cells. (**A**) 2D STED showing H2B (top row) or Flag-Net1 (middle row) immunofluorescence in G_1_and G_2_/M of the mitotic cell cycle, and quiescent cells. In top row, arrows denote rDNA structures. (**B**) High resolution 3D STED showing the different rDNA structures across the indicated cell states.

Although high-resolution microscopes are powerful for observing fine chromatin structures, it has been reported that the rDNA structure can look different depending on the orientation of yeast cells on slide glass if images are taken in two dimensions (2D)^23^. To circumvent this issue, we employed 3D STED microscopy (Fig. 1B). This technique allowed us to observe chromatin from different angles. By 3D STED microscope analyses, the rDNA exhibited pronounced nodes, which may represent “clutches” of either active or inactive copies of rDNA. The rDNA locus in G_1_ cells formed patch-like structures with multiple nodes. In G_2_/M cells, the rDNA was tightly condensed into its large loop structure with multiple rDNA nodes. In contrast, the quiescent rDNA occupies much smaller space than either G_1_ or G_2_/M cells, showing a high degree of compaction.

### Quiescent rDNA exhibits extensive *trans* chromatin contacts compared to G1 or G2/M cells

While microscopy provides a detailed view of both genomic and ribosomal DNA chromatin structures, we sought to further investigate the genomic interaction differences between the different cell cycle states. To this end, we leveraged existing Micro-C^25^ datasets from different stages of the mitotic cell cycle^22^ and quiescence^12^, focusing on the rDNA locus.

The rDNA locus consists of approximately 150 tandem repeats. Therefore, Micro-C data is represented as an average of all the rDNA copies shown over a single rDNA repeat. To obtain a larger-scale view of rDNA chromatin contacts, we first measured contact frequencies between the rDNA locus and the rest of the genome (Fig. 2A). To achieve this, we performed virtual 4C, an *in silico* method that quantifies the frequency of physical interactions between a chosen reference point, the rDNA locus, and other genomic regions. This analysis revealed stark differences in the relative frequencies of rDNA chromatin contacts between quiescent cells and cells in other cell-cycle stages. First, consistent with our 3D STED microscopy data, quiescent rDNA interacts more frequently with the rest of the genome, peaking at several sites in each chromosome. Second, in quiescence, the rDNA locus interacts much more frequently with the rest of chromosome XII compared to G1 and G2/M cells. Finally, contacts within the rDNA locus are much more frequent in quiescent cells.

**Figure 2.**
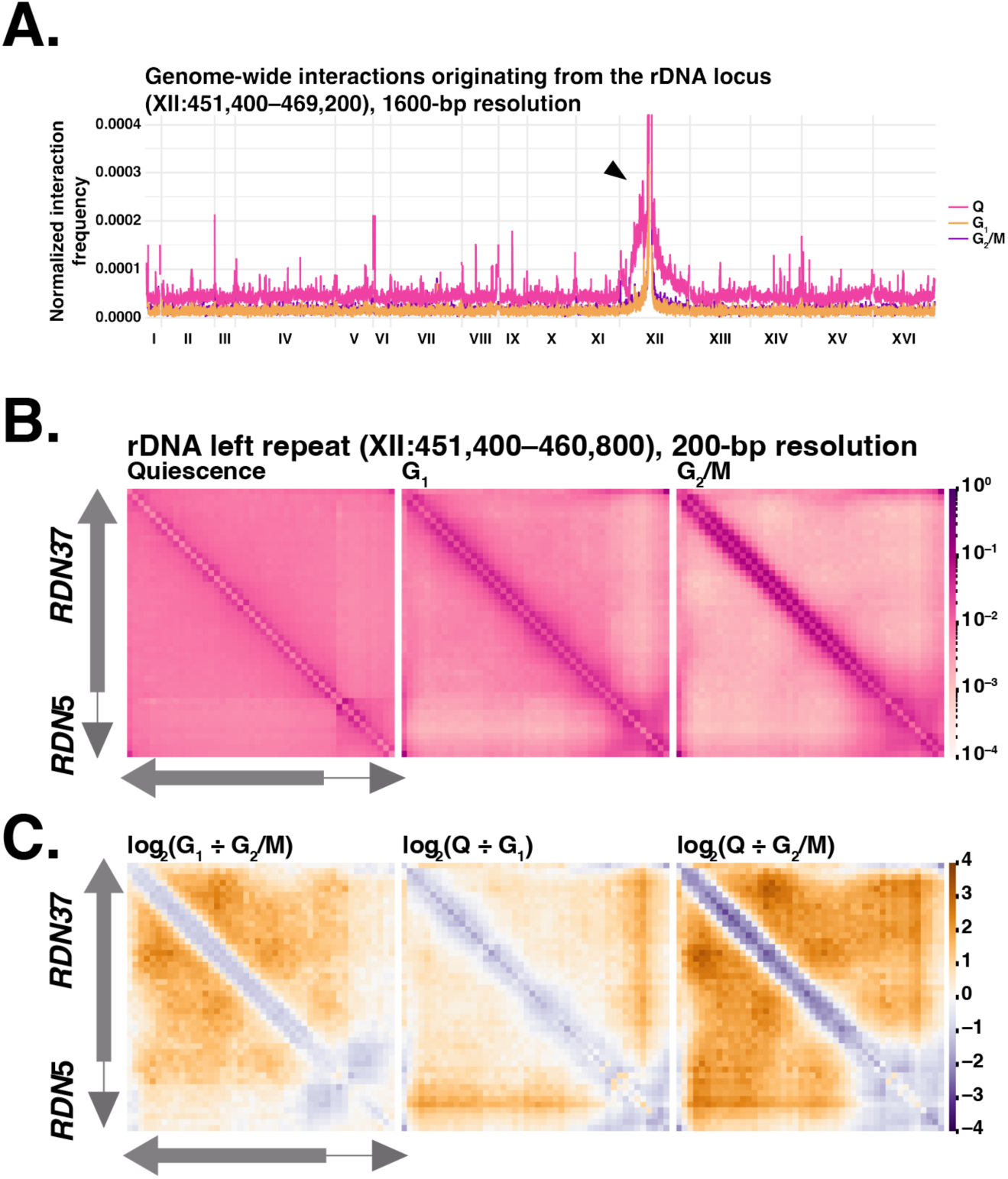
Micro-C analyses reveal genome-wide and local variations in rDNA chromatin contacts across different cell cycle stages. (**A**) Virtual 4C of normalized Micro-C interaction frequencies originating from the rDNA locus (arrow) for Q, G1-arrested, and G2/M-arrested cells. (**B**) Heatmaps showing averaged normalized interaction frequences across a single repeat unit of the rDNA locus in Q, G1-arrested, and G2/M-arrested cells. (**C**) Heatmaps displaying log_2_ ratios of normalized interaction frequences within the rDNA repeat, highlighting contact enrichment (gold) and depletions (purple) across pairwise combinations of cell cycle stages.

We next took a closer look at chromatin contacts within rDNA. The heatmaps in Fig. 2B show marked differences across the cell states. In G_2_/M, chromatin contacts along the diagonal line are highly prominent, demonstrating frequent short-range interactions. Additionally, contacts within the non-transcribed spacer (NTS), which contains 5S rDNA, a replication origin and a replication fork block, are relatively frequent. In contrast, rDNA chromatin in quiescent cells exhibits more medium-distance (off-diagonal) interactions. In addition, 35S rDNA and the NTS region form separate compartments and interactions between these compartments are relatively less frequent. Contact frequencies within the rDNA locus in G_1_ resemble an intermediate state between quiescence and G_2_/M. Pairwise comparisons of contact frequencies between cell-cycle stages support our conclusions (Fig. 2C).

### Sir2 has a quiescence-specific role in organizing the 3D structure of rDNA

Our microscopic and Micro-C analyses established that the rDNA locus forms quiescence-specific 3D chromatin structure that has not been described before. This means that the rDNA 3D structure represents the second large-scale, quiescence-specific 3D chromatin structure reported, with the first being telomere hyper-clustering^14,15^. In this phenomenon, all 32 telomeres in budding yeast nuclei coalesce into one or two large clusters during quiescence, in contrast to several (up to a dozen) smaller clusters in actively dividing cells cells^14,15^. Despite the striking formation of these structures in quiescence, the biological roles of both rDNA and telomere organization remain largely unknown.

To explore the biological roles of quiescence-specific large-scale 3D chromatin structures, we aimed to identify the factors responsible for the unique 3D chromatin organization of rDNA in quiescence. Because we could detect the RENT complex member Net1 on rDNA chromatin in quiescence, we reasoned that Sir2, another member of the RENT complex^24^, may be present on quiescent rDNA. Sir2 is an NAD-dependent histone deacetyl transferase that contributes to repressive chromatin structure at the rDNA locus, the mating-type loci and telomeres in actively dividing cells^26–30^. To evaluate the presence of Sir2 on the rDNA in quiescence, we performed chromatin immunoprecipitation followed by DNA sequencing (ChIP-seq) of Sir2. We found Sir2 is indeed bound at the rDNA locus during quiescence (Fig. 3A).

**Figure 3.**
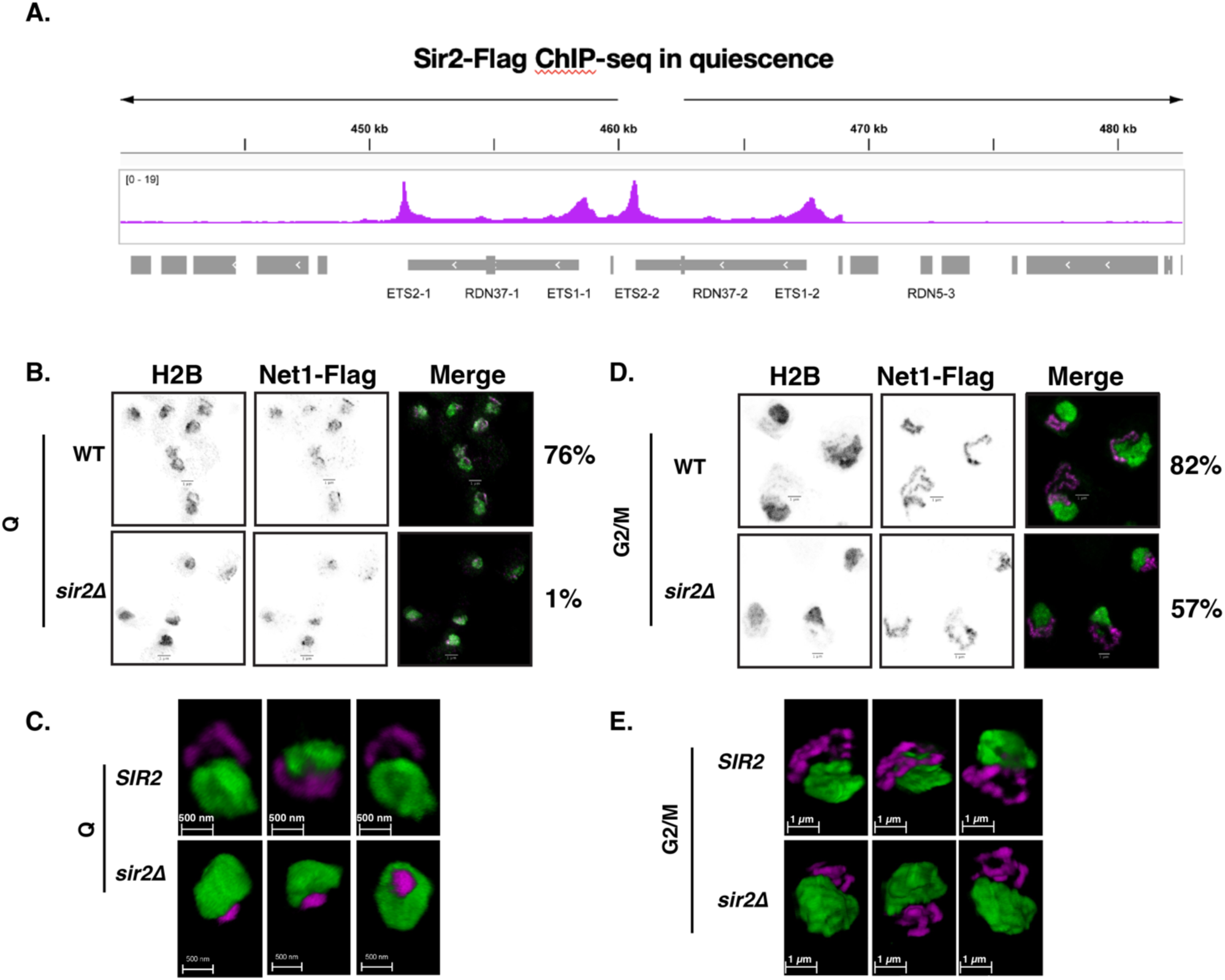
Sir2 is necessary for formation of quiescent-specific rDNA structure. (**A**) ChIP-seq of flag-tagged Sir2 in quiescent cells. Shown is the average of rDNA loci across two representative repeats. (**B**) 2D STED of quiescent cells with or without the *SIR2* gene. The numbers on the right denote the fractions of cells with condensed loop-like structure of rDNA. (**C**) 3D STED of quiescent cells as in (B). (**D**) 2D STED of G_2_/M cells with or without the *SIR2* gene. The numbers on the right denote the fractions of cells with rDNA loop structure. (**E**) 3D STED of G_2_/M cells as in (D).

To determine the consequence of deleting Sir2 on the quiescent rDNA, we employed STED microscopy in 2D (Fig. 3B and 3C). Because it has been established that Net1 binding to the rDNA locus is *SIR2*-independent^24^, we used Net1-FLAG to visualize the rDNA. In cells lacking the Sir2 protein, the rDNA structure in quiescence collapses into a small dot-like structure. (Fig. 3B, middle row). Indeed, using confocal microscopy to quantify the proportion of quiescent cells with loop-like structures, we found that these rDNA structures are mostly undetectable in *sir2* mutant cells (Fig. 3B, quantification on the right side of pictures). In contrast, rDNA loops in G_2_/M cells are visible in most *sir2* mutant cells, although the mutation slightly reduces the fraction of cells with the rDNA loop (Fig. 3D). These results showed that *SIR2* makes much larger contributions to rDNA 3D chromatin structure in quiescence than in G_2_/M. 3D STED analyses revealed more clearly the extent by which *SIR2* affects the rDNA 3D chromatin structure in quiescence (Fig. 3C) and G_2_/M (Fig. 3E).

### SIR proteins are essential for efficient quiescence entry and exit

With a mutant that disrupts quiescence-specific rDNA 3D chromatin structure in hand, we wondered what happens to quiescent-cell formation when rDNA structure is disrupted. We first noticed that the quiescent cell yield is much lower for *sir2* mutants compared to wild-type cells (Fig. 4A). The lower yield is due to two factors: First, mutants show a lower saturation density after a seven-day culture (Fig. 4A, pink); second, the fraction of quiescent cells among stationary-phase cells is significantly lower in mutants compared to wild-type cells (Fig. 4A, pink) Both findings indicate defects in long-term survival.

**Figure 4.**
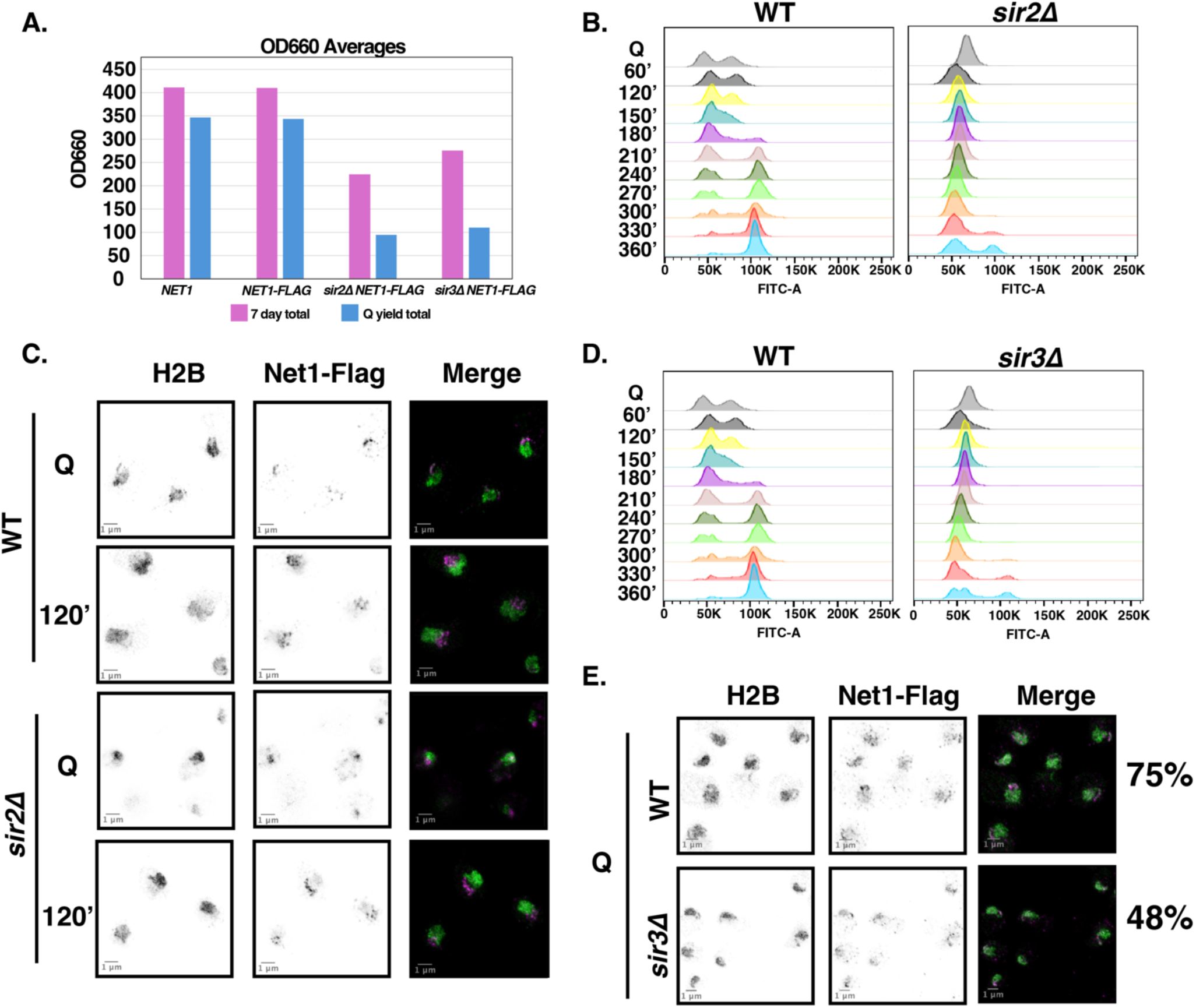
SIR proteins are necessary for proper quiescence release. (A) OD660 measurements of 7-day cultures (pink) and the quiescent fraction (blue). (B) Flow cytometry analysis of wild-type and *sir2*Δ cells released from quiescence (shown in minutes post-Q) using sytox to stain DNA shown by FIT-C channel. (C) 2D STED of cells released from quiescence with or without the *SIR2* gene. (D) Flow cytometrry analysis of wild-type and *sir3*Δ cells released from quiescence (shown in minutes post-Q) using sytox to stain DNA shown in by FIT-C channel. (E) 2D STED of quiescent cells with or without the *SIR3* gene. The numbers on the right denote the fractions of cells with condensed loop-like structure of rDNA.

Because quiescence is by definition a reversible G_0_ state^31^, we also sought to determine the effect of *SIR2* deletion on quiescence exit. To test this, we performed flow cytometry analysis of the cell cycle following release from quiescence (Fig. 4B). In cells lacking *SIR2*, we saw a notable ∼2.5-hour delay into the mitotic cell-cycle, suggesting that Sir2 is required for efficient release from quiescence. We also observed a difference in the quiescent peaks between wild-type and mutant cells, likely due to defects in the cell walls of Sir2-deficient cells. In wild-type cells, the characteristic double peak is attributed to the specialized cell wall formed during quiescence^32^.

Since we observed a dramatic quiescence exit defect in cells lacking *SIR2,* we wondered if this delay in quiescence release might also affect rDNA chromatin reorganization as cells re-enter the cell cycle. To test this, we used 2D STED microscopy to detect changes in the rDNA over time. By two hours, wild-type cells have fully formed the G_1_ “puff” (Fig. 4C). However, in mutant cells, the formation of a more wild-type-like quiescent rDNA structure only begins after around two hours. This suggests that the delayed rDNA reorganization is linked to the quiescent exit defects observed in *sir2* mutants. These results established that *SIR2* is required for both efficient quiescence entry and exit.

Deletion of *SIR2*, as well as *SIR3* and *SIR4*, has been reported to disrupt telomere hyper-clusters in quiescence^33,14,15^. Indeed, we found that *sir3* mutation causes quiescence entry (Fig. 4A) and exit (Fig 4D) defects to a very similar extent as that of the *sir2* mutation. However, distinguishing the functions of SIR proteins at telomeres versus. rDNA is challenging, as recent findings revealed that Sir3 is also present at rDNA and affects chromatin contacts^34^. In addition, Sir3-binding at the rDNA locus depends on *SIR2*’s deacetylase activity^34^, making it difficult to separate the effects of *sir* mutations at telomeres and rDNA. Consistent with this notion, we found that deleting *SIR3* significantly reduces the fraction of quiescent cells with loop-like structures, although the defect is less pronounced than that of *sir2* mutants (Fig 4E). Together, our data revealed that the disruption of two large-scale, quiescence-specific 3D chromatin structures leads to extensive defects in both quiescence entry and exit.

### Chromatin defects and Pol II mislocalization occur in quiescent cells lacking *SIR2*

While we have observed dramatic 3D chromatin defects in quiescent cells lacking the Sir2 protein, it remained unknown how the nucleosome landscape was impacted in cells lacking *SIR2*. To test this, we performed ChIP-seq of histone H3 and analyzed the rDNA locus (Fig. 5A). In quiescent cells lacking *SIR2*, H3 occupancy was higher at the rDNA locus compared to wild-type cells. We found that higher H3 occupancy at the rDNA locus is even more prominent in the first G_1_ phase upon quiescence exit (Fig. 5B), indicating that higher nucleosome occupancy at the rDNA locus persists through the process of quiescence exit.

**Figure 5.**
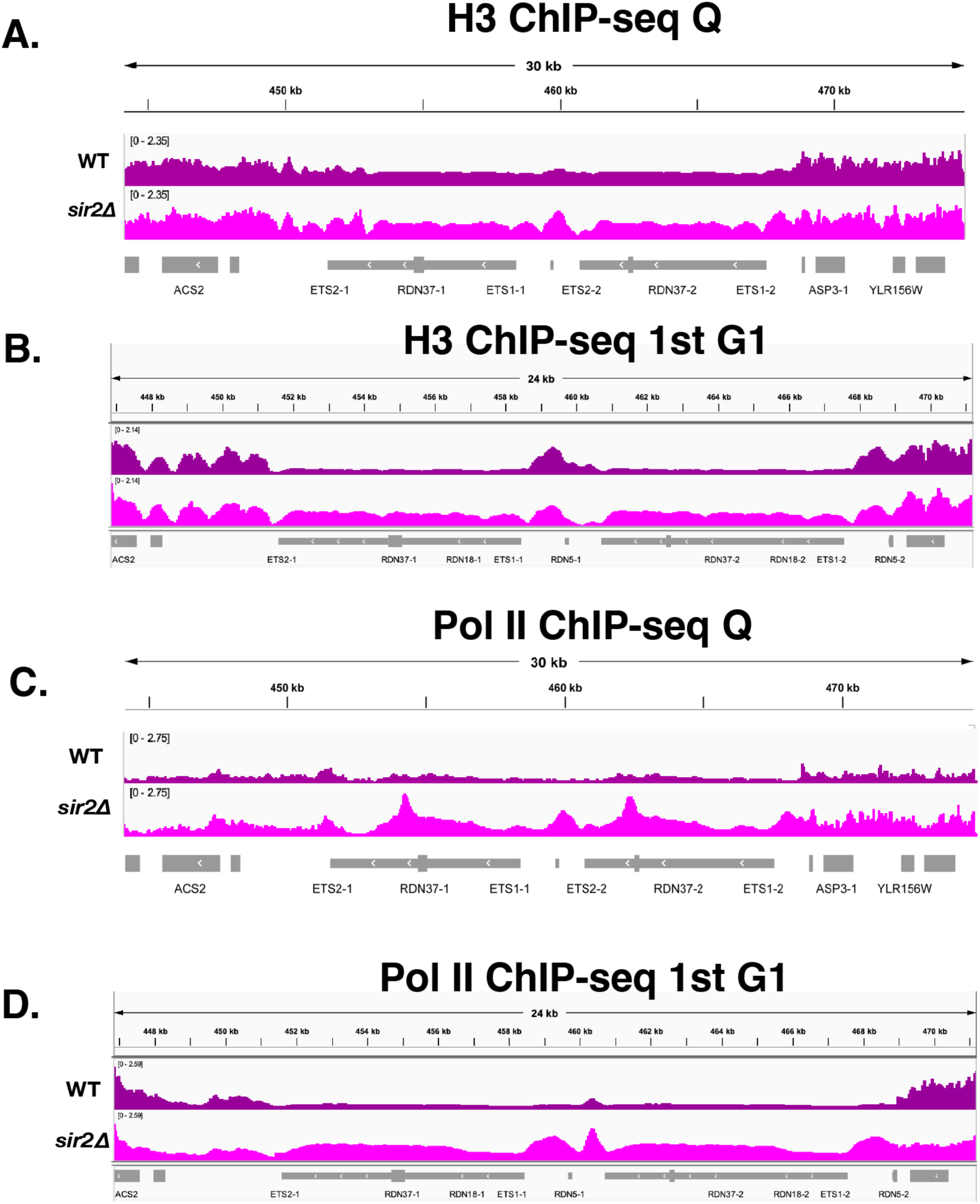
Chromatin structure and Pol II occupancy defects at the rDNA locus in quiescent cells lacking *SIR2*. (A) ChIP-seq of total H3 in quiescent cells. Shown is the rDNA locus, which is an average of ∼150 repeats represented in yeast by two copies of rDNA. Arrows denote NDRs seen in *sir2*Δ cells. (B) ChIP-seq of total H3 in the first G1 during quiescence exit. (C) Pol II ChIP-seq with and without *SIR2* in quiescence. (D) Pol II ChIP-seq with and without *SIR2* in quiescence during the first G1 upon quiescence exit.

While there was a notable increase across the rDNA locus in mutant cells, we observed two newly formed nucleosome-depleted regions (NDR) upstream and downstream of ETS-1 and ETS-2, respectively in quiescent cells (Fig. 5A). An aberrant NDR can often result in mislocalization of Pol II and, indeed, it has been shown that in the absence of Sir2, Pol II transcribes aberrant non-coding RNAs from the rDNA locus in actively dividing cells^35,36^. To test whether Pol II occupancy levels change in a Sir2-dependent fashion, we performed ChIP-seq of Pol II in quiescent cells. Wild-type cells had very little Pol II present at the rDNA locus, consistent with low levels of Pol II transcription across the genome as previously reported^7,12,37^ (Fig. 5C). In contrast, we observed elevated levels of Pol II present at the rDNA locus in quiescent *sir2* mutant cells, suggesting high levels of transcription by Pol II. Like defects in H3 levels, higher Pol II levels in *sir2* mutants persisted through quiescent exit, as higher levels of Pol II in the mutants were prominent in the first G_1_ phase out of quiescence (Fig. 5D). These results revealed that Sir2 lowers both nucleosome and Pol II occupancy at the rDNA locus in quiescence and throughout quiescence exit.

## DISCUSSION

Chromatin compaction is a well-established hallmark of quiescent cells, conserved across diverse organisms and cell types^8,11–13,38^. While the mechanisms controlling chromatin architecture in quiescence are still being elucidated, our previous work showed how smaller-scale structures, such as chromatin domains and folding of nucleosome arrays, are regulated through condensin and the H4 basic patch, respectively^12,13^. In contrast, this study reveals a larger-scale organization of 3D chromatin at the rDNA locus, suggesting a hierarchical regulation of chromatin structure during quiescence. We further demonstrated that this structure is regulated by the histone deacetylase Sir2 (Figure 6). Our findings provide evidence for the critical role of Sir2 in maintaining the rDNA structure during quiescence, contributing to both efficient quiescence entry and exit. We propose that Sir2 lowers nucleosome occupancy and represses aberrant long non-coding RNA (lncRNA) transcription by Pol II at the rDNA locus. One possible mechanism by which Sir2-mediated Pol II repression decreases nucleosome occupancy is through repression of transcription-coupled nucleosome assembly, which has been reported to take place upon transcription of long non-coding RNAs (lncRNAs) in yeast^39^. The most well-characterized is *SER3* gene repression by transcription of the lncRNA *SRG1* and nucleosome assembly^40^ (Figure 6). The disruption of this mechanism in *sir2* mutants leads to elevated nucleosome occupancy, aberrant rDNA folding, and defects in both quiescence entry and exit.

**Figure 6.**
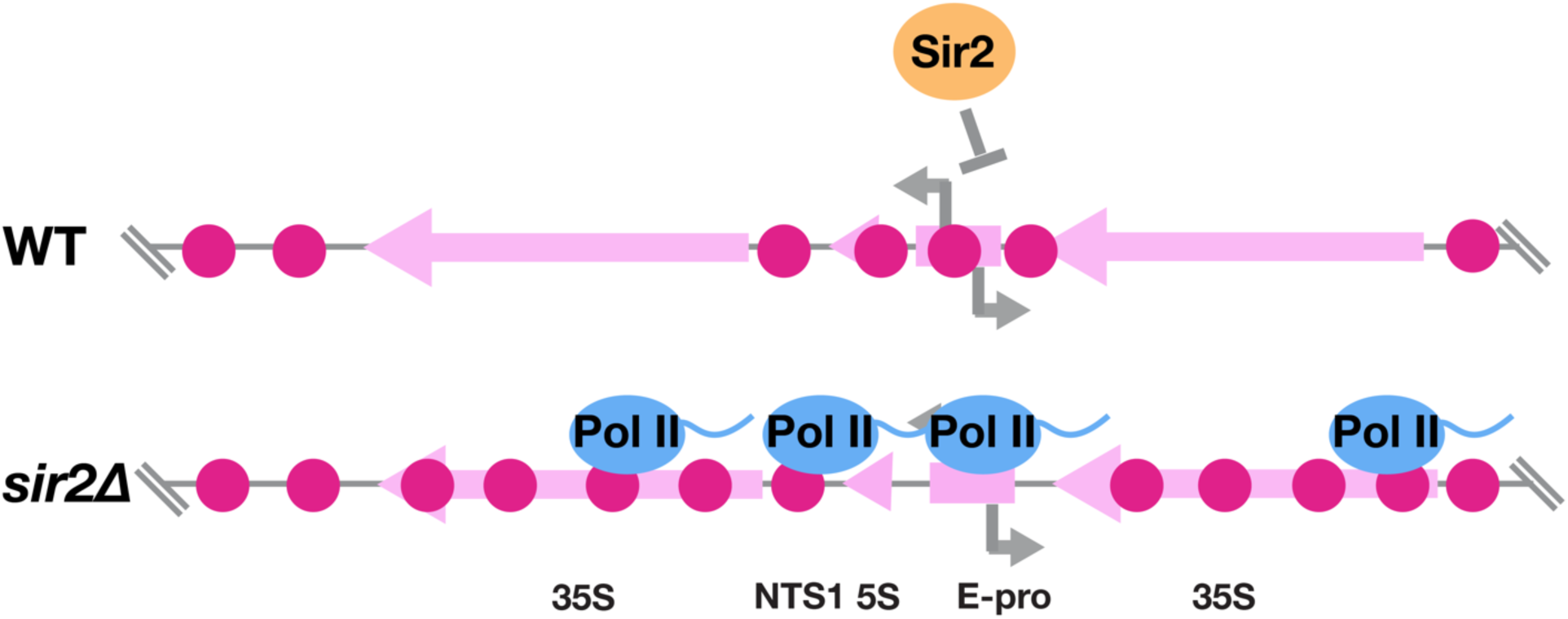
Sir2 is required for normal nucleosome occupancy and Pol II occlusion at the rDNA locus. In wild-type cells, the 35S rDNA gene is depleted of nucleosomes and intergenic regions have higher nucleosome occupancy. In cells lacking *SIR2* histone occupancy along the 35S gene is higher and nucleosome depleted regions form in intergenic loci. This chromatin disruption results in Pol II mislocalization to the rDNA locus.

Our discovery of the quiescent rDNA structure raises intriguing questions about its biological role. rDNA loops, described here, and telomere hyper-clustering, which was previously described by the Sagot and Taddei groups^14,15^, are required for efficient quiescence entry and exit. While the precise mechanism remains to be elucidated, these large-scale chromatin structures may serve as organizational hubs, facilitating the rapid reactivation of genes necessary for cell cycle re-entry^37^. These structures may also help maintain the genomic integrity of highly repetitive regions like rDNA, where transcriptional misregulation could lead to genome instability^41^.

Sir2’s role in regulating rDNA structure adds to its well-documented function in chromatin silencing at multiple genomic loci, including telomeres and mating-type loci^26–29^. Our results align with previous studies showing that Sir2 is involved in suppressing aberrant lncRNA transcription from the rDNA locus^30,35,36^. By preventing lncRNA transcription, Sir2 likely reduces nucleosome occupancy and maintains a specific chromatin configuration in quiescence. The increased nucleosome occupancy and 3D chromatin misfolding observed in *sir2* mutants support this model, suggesting the possibility that transcription-coupled nucleosome assembly may play a key role in regulating rDNA structure.

Given the parallels between rDNA organization and other quiescence-specific chromatin structures, it is possible that Sir2 has a broader role in modulating chromatin architecture during quiescence, and similar mechanisms may be conserved in mammals involving Sir2 homologs such as SIRT1^42^. The observed defects in quiescence exit in *sir2* mutants may reflect a more general failure to establish the chromatin configurations required for rapid cell cycle reactivation. The increased Pol II mislocalization we observed in *sir2* mutants further supports this hypothesis, as proper chromatin organization is essential for accurate transcriptional regulation during quiescence exit^37^.

Beyond yeast, our findings have potential implications for understanding quiescence in metazoans. Quiescent cells in multicellular organisms, such as those with stem cells, must similarly maintain genome stability and transcriptional repression during periods of cell cycle withdrawal. The role of chromatin structure in maintaining the quiescent state is becoming increasingly evident, and it is possible that similar mechanisms, involving Sir2 homologs or other chromatin regulators, contribute to the regulation of quiescence-specific chromatin structures in mammals. Investigating whether these findings are conserved in higher organisms could provide valuable insights into stem cell biology and cancer, where improper regulation of quiescence is linked to disease progression.

Together, our work highlights a previously unrecognized role for Sir2 in organizing quiescence-specific chromatin structures at the rDNA locus. These results not only expand our understanding of large-scale 3D-chromatin regulation in quiescence but also provide a framework for future studies exploring the broader significance of large-scale chromatin structures in cell state transitions. Further investigation into the factors governing these structures, as well as their potential conservation in other systems, will be critical for advancing our knowledge of quiescence and its impact on cell and organismal survival.

## METHODS

### Yeast strains and growth conditions

The *Saccharomyces cerevisiae* strains used in this study are isogenic to the W303-1a strain, with a corrected *rad5* allele as described^43^. A detailed list of strains is provided in the strain table (Supplementary Table 2) In general, we generated new yeast strains through yeast homologous recombination using DNA primers containing 50-bp of overlap to the region of interest and a deletion or insertion cassette as described in^44^. All log cell cultures were grown in YPD medium (2% bacto peptone, 1% yeast extract, 2% glucose).

### Alpha-factor mediated G1 arrest

Cells were grown to early log phase and then incubated with 5ug/ml alpha factor for 90 minutes or less, depending on visual inspection of the culture and percentage of cells with the ‘shmoo’ morphology (∼90% by eye)^45^. G_1_ arrest was verified using flow cytometry to measure total DNA content of the cells (described below). Because *sir2Δ* mutant cells have mating defects due to ineffective silencing of the *HML* loci^26^, we used a strain containing a deletion of *HML1* to generate alpha-factor responsive cells.

### Quiescent cell induction and collection

To generate quiescent cells, we inoculated YPD cultures and incubated the cells at 30°C. It was important to adjust the pH of the YPD medium of 5.5 using HCl, as described^46^. Quiescent cell cultures were grown with a medium-to-flask volume ration of 1:10 to ensure optimal aeration for the development of healthy quiescent cells. Cells were incubated at 180 RPM on an orbital shaker.

After seven days of growth, cultures were pelleted, rinsed with double-distilled H2O (ddH2O), resuspended in 1 mL ddH2O, and gently layered onto a pre-prepared percoll gradient. For each gradient, 400 optical density (OD)_660_ of cells were applied to a 25 mL gradient tube. Gradients were centrifuged at 1,000 RPM for 1 hour at 4°C. Following centrifugation, the upper layer of non-quiescent cells and the middle ∼8 mL layer were carefully removed by pipetting. The quiescent cell layer was washed twice with ddH2O in a 50 mL conical tube at 3,000 RPM for 10 minutes per wash.

For quiescence release experiments, cells were added to flasks containing YPD pH 5.5 as described above with 2% glucose to a final concentration of 0.5 optical density units ODU/mL. Cells were incubated at 25°C, shaking at 180 RPM for the time indicated before harvest.

### Flow cytometry to measure DNA content and infer cell cycle stages

To prepare cells for flow cytometry, 0.5 mL of log-phase culture at OD₆₆₀ equal to 0.5 or 0.25 ODU of purified quiescent cells was fixed in 70% ethanol and stored at 4°C overnight. The fixed cells were then washed with water and resuspended in 0.2 mL of 2 mg/mL RNase A solution, followed by incubation for 4 hours at 37°C. After RNase A treatment, cells were pelleted, resuspended in 0.2 mL of 2 mg/mL Proteinase K, and incubated for 45 minutes at 50°C.

Following Proteinase K treatment, cells were pelleted again and resuspended in 0.2 mL of 50 mM Tris-HCl, pH 7.5, and incubated overnight at 4°C. For flow cytometry analysis, cells were briefly sonicated, and an aliquot (50–100 µL) was combined with 1 mL of a 1x SYTOX Green solution (50 mM Tris-HCl, pH 7.5; Fisher Scientific, catalog #S7020). After incubation at room temperature for 1 hour in the dark, samples were analyzed on a Becton Dickinson Canto II cytometer, and the data were processed using FlowJo software.

### Fluorescence microscopy

G1 cells were arrested with 5 μg/ml α-factor as described above. G2/M cells were arrested with 10 μg/ml Nocodazole. Q cells were purified as described above. 0.5 ODU of A660 nm were saved for G1 and G2/M, while 10 units were saved for Q cells. Cells were fixed in 3.7% formaldehyde solution in 0.1M KPi for 25 minutes at room temperature. Samples were washed with 0.1M KPi and then 1.2M sorbitol-citrate. Cells were digested with zymolyase 100T at 10 μg/ml for G1 and G2/M cells, and 0.5 mg/ml for Q cells, in a solution with 0.5% β-mercaptoethanol and 1.2M sorbitol-citrate. Digestion was done at 30°C with 600 RPM shaking, until approximately 50–70% of cells had been spheroplasted, as judged by spectrophotometer measurement of optical density. Cells were washed and then resuspended in 1.2M sorbitol-citrate for incubation on slides coated with 0.1% polylysine. The slides were washed with ice cold methanol for three minutes, followed by ice cold acetone for 10 seconds. Once dry, the slides were washed once with PBS-BSA, then incubated with PBS-BSA for at least one hour at room temperature. Samples were incubated with PBS-BSA containing the primary antibodies, at 1:100 dilution of α-FLAG antibody and 1:500 to 1:4000 dilution of the α-H2B antibody for one hour to overnight, depending on the type of microscopy they would be used for. The samples were washed four times for five minutes each with PBS-BSA. The samples were incubated with PBS-BSA containing the secondary antibodies at 1:200 dilution for ATTO 647N and 1:200 dilution for Alexa 594 for a minimum of one hour or maximum of overnight. The samples were washed four times for five minutes each with PBS-BSA. Samples were mounted with ProLong Gold antifade mountant and incubated at room temperature for at least 24 hours before imaging. Samples were imaged within two weeks of preparation.

Samples were imaged on a Leica SP8 confocal microscope with 2D STED capabilities, or a Leica Stellaris 8 confocal microscope with 3D STED capabilities. All images were taken using Leica Application Suite X (LAS X) software, version 3.5.7 (for 2D STED and *sir2*Δ confocal imaging) or 4.7.0 (3D STED and *sir3*Δ confocal imaging). Z-stacks were taken for each field with total stack size varying between 1.82 – 7.76 µm. Step size was constant for each imaging modality, where confocal images used 0.3 μm, 2D STED used 0.156 μm, and 3D STED used 0.065 μm between steps. All imaging was done with a white light excitation laser and detection on HyD, HyD S or HyD X spectral detectors. 3D STED images were acquired on HyD X detectors operating in TauSTED mode. The excitation wavelengths and emission detection ranges were set to minimize crosstalk between Alexa594 (labeling FLAG) and Atto647N (labeling H2B). 775 nm STED depletion was used for both dyes. Confocal imaging used a HC PL APO CS2 63x/1.4 oil objective; 2D STED imaging used a HC PL APO CS2 100x/1.4 oil objective, and 3D STED imaging used a HC PL APO CS2 STED 93x/1.3 glycerol objective with motorized correction collar.

All 2D STED and confocal images were deconvolved with Leica Lightning, then maximum intensity projections of the z-stacks were generated and scale bars were added using FIJI / ImageJ (version 2.9.0 / 1.53u)^47^. These processed confocal images were blinded for genotype and used to quantify the frequency of rDNA loops by manually assessing each cell for the presence or absence of a loop. 3D STED images and videos were rendered using Leica LAS X 3D image viewer (version 3.5.7), with rotation around the y-axis.

### ChIP-seq

Chromatin immunoprecipitation (ChIP) was performed in biological duplicate using isolates from two different parent strains and using a modified protocol based on Rodriguez et al., 2014^48^. A total of 200 OD₆₆₀ units of cells were crosslinked and sonicated. For each reaction, proteins were immunoprecipitated from 1 µg of chromatin using 1 µL anti-H3 antibody (Abcam, 1791), conjugated to 20 µL of protein G magnetic beads (Invitrogen, 10004D). Pol II ChIPs were conducted with an antibody targeting the Rpb1 subunit (10 µL per reaction, Cell Signaling 2629S), similarly conjugated to 20 µL of protein G magnetic beads. For Sir2 ChIPs, a Flag M2 mouse monoclonal antibody (Sigma-Aldrich, F1804) was used, targeting the Flag epitope tag and conjugated to protein G beads. Library preparation was carried out using the Ovation Ultralow v2 kit (Tecan, 0344), and libraries were sequenced as 50 bp paired end reads on an Illumina NextSeq platform. Sequencing was conducted at the Fred Hutchinson Cancer Center Genomics Facility.

### ChIP-seq Analysis

Raw reads were aligned to the sacCer3 reference genome using Bowtie2^49^. The aligned reads were then filtered with SAMtools^50^. Input-normalized ChIP-seq data were generated as bigwig files from the filtered BAM files using deepTools2^51^, normalizing the immunoprecipitation (IP) data by dividing it by the input data. All ChIP-seq IP data were normalized to RPKM and the corresponding input samples. Coverage files were visualized in the Integrated Genome Viewer (IGV)^52^.

## Micro-C Data Processing and Analysis

### Trimming and aligning Micro-C sequenced reads

Using the program Atria (version 3.2.2)^53^, raw .fastq files underwent adapter and quality trimming to remove low-quality sequences and adapters. The trimmed reads were then aligned against the sacCer3 reference genome using BWA MEM (version 0.7.17)^54,55^. Post-alignment, the reads were compressed into .bam format using Samtools (version 1.16.1)^50^.

### Parsing and Filtering rDNA Contacts

Micro-C data preprocessing began by parsing .bam files into contact pairs using Pairtools^56^ (v1.0.2), focusing on both genomic contacts and specific contacts originating from the rDNA locus on chromosome XII. For genomic contacts, a minimum MAPQ score of 1 was used to exclude multi-mapping reads, while rDNA contacts were parsed with a MAPQ score of 0 to accommodate the repeated rDNA structure in the *S. cerevisiae* genome. Following parsing, sorted pairs were deduplicated to remove PCR/optical duplicates, and rDNA contacts were isolated using filtering criteria that excluded contacts outside the rDNA locus.

### Contact Matrix Construction and Normalization

Contact matrices were constructed from parsed data using the Cooler package (v0.9.2)^57^, binning the genome at 25-bp intervals. For visualization, the Cooler zoomify program generated multi-resolution matrices, ranging from 50 bp to 102.4 kb, allowing for multi-scale analysis. Normalization was performed with the Knight-Ruiz algorithm to correct for biases^58^, with a maximum of 2,000 iterations specified for balancing.

### Downsampling and Conversion for Visualization

To ensure comparable matrix sums across different cell states (quiescent, G1, G2/M), a custom Python script was used to randomly downsample matrices to the smallest contact sum among the samples. These matrices were balanced and converted to .hic format using the FAN-C package (version 0.9.27)^59^, enabling compatibility with HiGlass and other visualization tools.

### Analysis of rDNA-Specific Contact Patterns

For focused analyses on the rDNA left array, FAN-C was used to subset data corresponding to the region on chromosome XII, positions 451,400–460,800. These rDNA-specific matrices were then visualized using .cool-formatted files.

### Comparative and Visualization Analysis

Pairwise comparisons between cell-state-specific contact matrices were performed using HiCExplorer’s^60^ hicCompareMatrices tool. Contact matrices were visualized using hicPlotMatrix with log transformations where applicable. Code for Micro-C data processing and analyses are deposited in the following repository: <u>github.com/kalavattam/2024_Sir2</u>.

### Virtual 4C Analysis of rDNA Interactions

Virtual 4C (v4C) analyses were conducted to examine interactions between the rDNA locus and other genomic regions. Using HiCExplorer’s hicPlotViewpoint^60^, virtual contacts originating from the rDNA region were computed with respect to *cis* and *trans* chromosome regions, producing state-specific genome-wide interaction profiles. Resulting data were further analyzed and visualized in R.

## Acknowledgements

We are grateful to Linda Breeden and Shawna Miles for sharing critical insights on *sir2Δ*.We thank Lena Schroeder and Hoku West-Foyle for STED microscope training as well as feedback on the manuscript, Susan Parkhurst and Mitsutoshi Nakamura for critical advice on microscopy, Ilya Flyamer and Seungsoo Kim for discussion on Micro-C analyses, and the Tsukiyama lab members for comments on the manuscript. TT was supported by R35GM139429. This research was supported by the Cellular Imaging Shared Resource, <u>RRID:SCR_022609</u>; the Flow Cytometry Shared Resource, <u>RRID:SCR_022613</u>; and the Genomics & Bioinformatics Shared Resource, <u>RRID:SCR_022606</u>, of the Fred Hutch/University of Washington/Seattle Children’s Cancer Consortium (P30 CA015704).

## FIGURES & LEGENDS

Supplemental Table 1.

Supplemental Table 1 **G2/M**
| **Strain** | **rDNA loop detected** | **Total cells counted** | **Percent rDNA structure detectable** |
| --- | --- | --- | --- |
| YTT7766 G2/M | 234 | 284 | 82% |
| YTT7786 G2/M | 178 | 313 | 57% |

**G2/M**
| **Strain** | **rDNA loop detected** | **Total cells counted** | **Percent rDNA structure detectable** |
| --- | --- | --- | --- |
| YTT7767 G2/M | 216 | 256 | 84% |
| YTT7787 G2/M | 157 | 255 | 62% |

**Quiescence**
| **Strain** | **rDNA loop detected** | **Total cells counted** | **Percent rDNA structure detectable** |
| --- | --- | --- | --- |
| YTT 7766 Q | 185 | 280 | 66% |
| YTT 7786 Q | 9 | 295 | 3% |

**Quiescence**
| **Strain** | **rDNA loop detected** | **Total cells counted** | **Percent rDNA structure detectable** |
| --- | --- | --- | --- |
| YTT 7767 Q | 321 | 421 | 76% |
| YTT 7787 Q | 7 | 524 | 1% |

**Quiescence**
| **Strain** | **rDNA loop detected** | **Total cells counted** | **Percent rDNA structure detectable** |
| --- | --- | --- | --- |
| YTT 7766 Q | 179 | 255 | 70% |
| YTT 7833 Q | 254 | 523 | 49% |

**Quiescence**
| **Strain** | **rDNA loop detected** | **Total cells counted** | **Percent rDNA structure detectable** |
| --- | --- | --- | --- |
| YTT 7767 Q | 237 | 300 | 79% |
| YTT 7834 Q | 198 | 416 | 48% |

**Supplemental Table 2.** Yeast strains used in this study.

| Strain | Genotype |
| --- | --- |
| YTT5781 | *MATa RAD5+* |
| YTT5783 | *MATa RAD5+* |
| YTT7766 | *MAT a RAD5+ can1-100 NET1-2L3FLAG-kanMx* |
| YTT7767 | *MAT a RAD5+ can1-100 NET1-2L3FLAG-kanMx* |
| YTT7486 | *MAT a RAD5+ can1-100 sir2::Hyg Net1-2L-3FLAG-NatMx* |
| YTT7487 | *MAT a RAD5+ can1-100 sir2::Hyg Net1-2L-3FLAG-NatMx* |
| YTT7450 | *MATa can1-100 RAD5+ sir2::HygMx* |
| YTT7451 | *MATa can1-100 RAD5+ sir2:: HygMx* |
| YTT7488 | *MATa can1-100 RAD5+ sir2::Hyg hml::KanMX* |
| YTT7489 | *MATa can1-100 RAD5+ sir2::Hyg hml::KanMX* |
| YTT7566 | *Mat a prototroph Sir2-2L3FLAG- HygMx* |
| YTT7567 | *Mat a prototroph Sir2-2L3FLAG- HygMx* |

